# Developmental switch in prediction and adaptation to pain in human neonates

**DOI:** 10.1101/2022.04.05.486988

**Authors:** Mohammed Rupawala, Oana Bucsea, Maria Pureza Laudiano-Dray, Kimberley Whitehead, Judith Meek, Maria Fitzgerald, Sofia Olhede, Laura Jones, Lorenzo Fabrizi

**Author notes:** These authors should be considered as joint senior authors (Laura Jones, Lorenzo Fabrizi). **Funding Information**, This work was funded by the Medical Research Council UK (MR/S003207/1) and the European Research Council (CoG 2015-682172NETS) within the Seventh European Union Framework Program. OB was supported by the Canadian Institutes of Health Research (FBD–170829). **Authors’ Contributions** Conceptualisation of study: L.J., L.F., M.F.; data collection and preparation: L.J., M.P.L-D., K.W.; clinical supervision: J.M.; data analysis: M.R., L.J., O.B., S.O.; data interpretation: M.R., L.J., L.F.; manuscript preparation: M.R., L.J., L.F. All authors discussed the results and commented on the manuscript.

## Abstract

Habituation to recurrent non-threatening or unavoidable noxious stimuli is an important aspect of adaptation to pain and indicates the ability of the brain to encode expectation of imminent nociception. However, it is not known whether the newborn brain can predict and habituate to recurrent noxious inputs. We used electroencephalography to investigate changes in cortical microstates, which represent the complex sequential processing of noxious inputs, following repeated clinically-required heel lances in term and preterm infants. Noxious stimulus repetition decreased the engagement of early sensory-related microstates and associated behavioural and physiological responses in term infants, while preterm infants did not show signs of adaptation. Nevertheless, both groups displayed a switch between different microstates at longer latencies. These data suggests that the preterm brain is capable of encoding high-level contextual differences in pain, but cannot update its prediction, which allows for adaptation, emphasising the vulnerability of this population to recurrent pain.

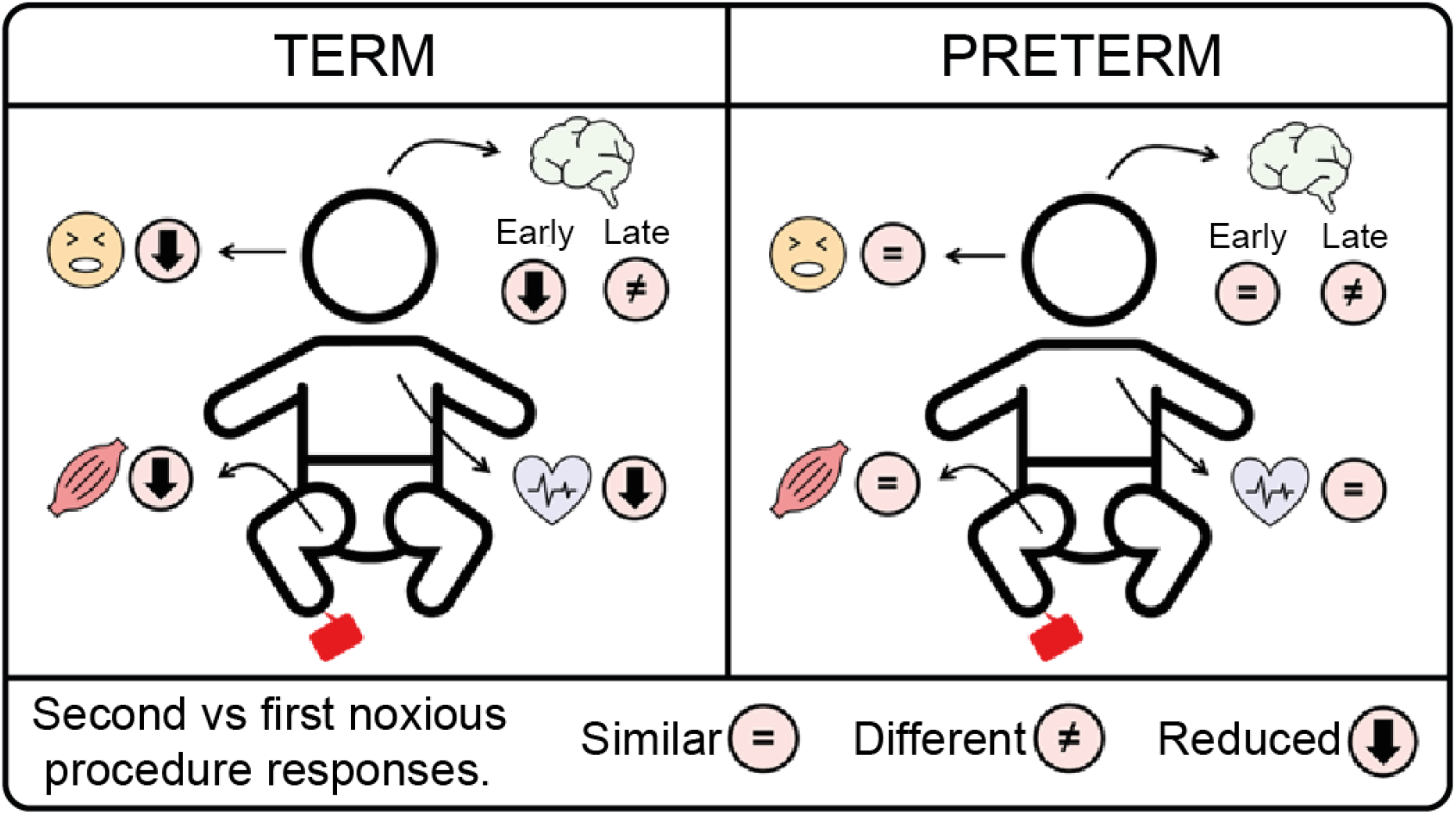

## INTRODUCTION

Habituation, described as decreased responses to repeated stimulation, is the simplest manifestation of behavioural plasticity and is considered a basic form of memory/learning (Groves & Thompson, 1970). Pain is a multifaceted experience which requires extensive cortical processing that ultimately results in the appropriate behavioural and physiological responses ensuring physical integrity and survival. Pain habituation is therefore an important physiological and psychological adaptation to recurrent or sustained noxious stimuli and is likely to preserve physical, emotional, and cognitive resources in favour of more pressing goals when the threat is unavoidable or not life-threatening (De Paepe et al., 2019).

Habituating cortical activity to repeated stimuli has been observed across a range of stimulus properties (e.g., frequency, duration, position), time scales, sensory modalities, and brain areas (Grill-Spector et al., 2006). In the healthy adult brain, pain habituation is associated with modulation of parallel and sequential pathways (Kucyi & Davis, 2015) including decreased activity in areas related to pain sensation, such as the thalamus, insula and primary (SI) and secondary (SII) somatosensory cortices and increased activity in areas involved in descending pain modulation, such as the anterior cingulate cortex (ACC) (Bingel et al., 2007). These areas encode the sensory-discriminative, affective-motivational, and cognitive-evaluative components of the pain experience and initiate the relevant top-down signal processes that execute physical behaviours to avoid pain chronicity and resolve physical injury. These cortical processes also engage descending opioidergic mechanisms in the periaqueductal gray (PAG) projecting to the rostral ventromedial medulla (RVM) resulting in the modulation of spinal reflex response (Bromm & Scharein, 1982), heart rate changes (Colloca et al., 2006) and skin conductance response (Bromm & Scharein, 1982).

Modern concepts of perception propose that these cortical activity changes arise from the integration of sensory input with predictions about the expected stimulus based on a continuously updated internal model of the world (Friston & Kiebel, 2009). Any differences between the expected and received input would therefore lead to a prediction error reflected in the activation of brain areas known to be involved in pain processing (Geuter et al., 2017; Wang et al., 2022). With respect to pain habituation, this concept suggests that sensory brain responses would be reduced if the brain was able to compare and successfully predict the arrival of successive unavoidable or non-threatening stimuli (Seymour & Mancini, 2020).

This type of learning and prediction is likely to evolve over the lifespan as the brain and the underlying connections develop in conjunction with exposure to a wide range of sensory experiences. However, at what stage in human development such complex cortical processes manifest and how effectively they enable modulation of physiological responses, remains unknown. The brain goes through very rapid maturational changes over the perinatal period (Kostović & Jovanov-Milošević, 2006) while hospitalised neonates, including those who are sick and/or born preterm, are exposed to several noxious procedures every day as part of their clinical care (Laudiano-Dray et al., 2020). Here we hypothesise that the ability of the brain to adapt to repeated noxious stimuli and to drive appropriate behavioural and physiological adaptive responses, to balance threat avoidance and energy wastage, changes over the perinatal period.

A heel lance is the most frequently performed blood sampling test in hospitals for newborns and occasionally, albeit rarely, has to be repeated in quick succession to collect the required amount of blood (Shah & Ohlsson, 2011). Each lance elicits strong behavioural, physiological, spinal and specific cortical responses (Cornelissen et al., 2013; Jones et al., 2017; Slater, Cornelissen, et al., 2010; Slater, Worley, et al., 2010). Here we present cortical, autonomic and somatic responses from human term and preterm neonates that required two consecutive and identical clinical heel lances and demonstrate distinct developmental changes in the adaptation and prediction of repeated unavoidable noxious stimuli.

## RESULTS

Twenty-one human infants (table 1, methods), of which 10 preterm (median 34 completed postmenstrual weeks, range 32-36 weeks), and 11 term equivalent (which includes four preterm-born and seven term-born neonates that were studied at term age, henceforth referred to as ‘term’, median 39 completed postmenstrual weeks, range 37-44 weeks) underwent two blood tests (heel lances) in brief succession (3-18 minutes separation, as clinically required), in the same heel skin area (figure 1). The two groups did not differ in postnatal age (preterm mean (SD): 26.4 (23.09) days, term mean (SD): 24.45 (34.04) days, t(19)=0.15, p=0.88, 95% CI = −24.91 – 28.80) nor in interstimulus interval (preterm mean (SD): 8.91 (3.82) mins, term mean (SD): 6.96 (2.39) mins, t(19)=1.42, p=0.17, 95% CI = −0.92 – 4.89), using an independent samples Student’s t-test. Moreover, there was no significant change in sleep state distribution across subjects between first and second lance (figure 6, methods) and all infants (except one) stayed in the same position (cot, skin-to-skin or held) for both lances (i.e., sleep and position were not covariates between repeated lances).

**Table 1:** Infant demographics.

| | Preterm<br>($< 37$ weeks PMA at study) | Term<br>($\geq 37$ weeks PMA at study) |
| --- | --- | --- |
| *N* | 10 | 11 |
| *GA at birth (weeks<sup>+</sup>days)* | 31<sup>+6</sup> (24<sup>+2</sup> – 34<sup>+5</sup>) | 38<sup>+0</sup> (24<sup>+6</sup> – 41<sup>+4</sup>) |
| *PNA (complete days)* | 14 (5 – 65) | 13 (1 – 116) |
| *PMA at study (weeks<sup>+</sup>days)* | 34<sup>+0</sup> (32<sup>+6</sup> – 36<sup>+4</sup>) | 39<sup>+6</sup> (37<sup>+4</sup> – 44<sup>+2</sup>) |
| *No. female* | 5 (50%) | 4 (36%) |
| *Birth weight (g)* | 1461 (708 – 2240) | 3352 (600 – 4070) |
| *No. caesarean deliveries* | 7 (70%) | 6 (55%) |
| *No. stimulation on right foot* | 2 (20%) | 7 (64%) |

Note: Gestational age (GA) is the number of weeks (and additional days) elapsed from the first day of the mother's last menstrual cycle to the day of delivery. Postnatal age (PNA) is the number of days elapsed since birth. Postmenstrual age (PMA) is the sum of these two ages. Values represent median and range.

**Figure 1:**
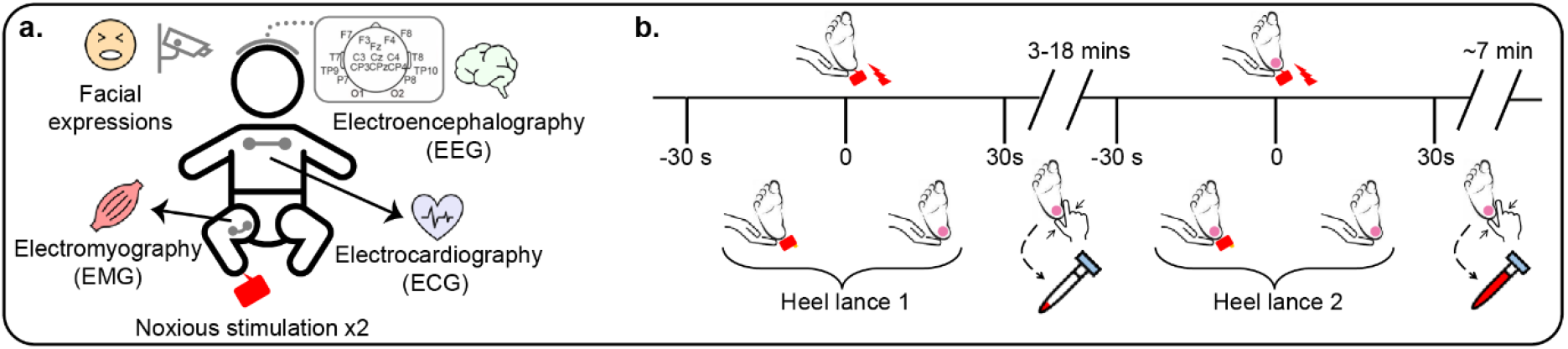
a) Recording setup. b) Timeline of events. All recordings began before the first heel lance and continued for the entire duration of the study. In the 30 s preceding each lance attempt (i.e., between −30 – 0 s) the lancet was held against the heel. The lance was released at time 0 s and squeezing started only 30 s after to obtain a post-lance response free from other stimuli. As not enough blood was collected after the first sampling attempt, the same procedure was repeated after 3-18 minutes.

Changes in the cortical, autonomic, and somatic (brainstem and spinal levels) activity of the neonatal nervous system were recorded simultaneously following each skin-breaking procedure as follows: 1) cortical activity using scalp electroencephalography (EEG), 2) reflex flexion withdrawal of the stimulated limb using biceps femoris electromyography (EMG), 3) heart rate using electrocardiography (ECG) and 4) facial expression behaviour from video recording (figure 1).

### A heel lance engages a distinct sequence of cortical microstates in term and preterm infants

We first investigated the effect of heel lance repetition upon single channel vertex (Cz) event-related potentials (ERPs). These were significantly modulated in term, but not preterm infants (figures 2a and 3a, supplementary figure 1). In term infants, differences in activity consisted of a reduction in positive voltage up to 78 ms following the second lance compared to the first and to a complete reverse in signal polarity between 599-734 ms from negative to positive.

**Figure 2:**
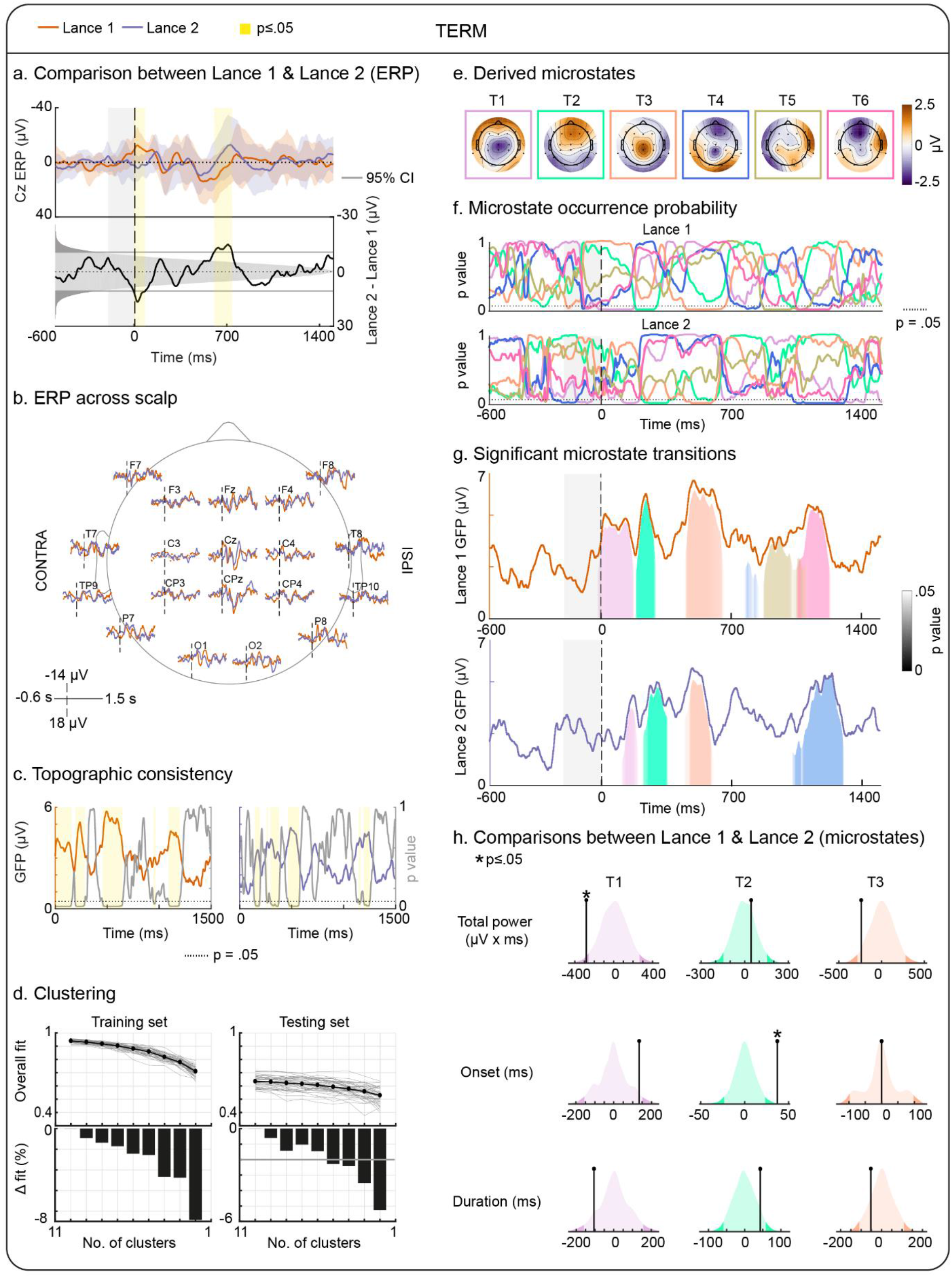
Term neonate global event-related field topography (microstate) analysis of EEG data (n=11) to repeated lances. a) ERPs at the vertex (Cz) in response to first and second lance. Top panel: mean (orange and purple lines) and standard deviation (corresponding shaded areas) across subjects. Bottom panel: difference between mean responses to first and second lance, non-parametric distribution of differences between surrogate mean activity (shaded grey), and values below the 2.5^th^ and above the 97.5^th^ percentiles (thick grey regions and lines). Yellow areas represent time-points with significant differences between first and second lance. b) Group average ERPs from each of the 19 channels on the infants ‘ scalp. c) Topographic consistency test results for first and second lance. Note how a high GFP corresponds to a lower p-value (grey) as this represents a strong spatial pattern consistently present across infants in the group. d) Cross validation testing to determine the optimal microstate basis dimension. A random 50% of subjects were used to derive clusters (i.e., microstates) and the remaining used for microstate fitting in each iteration. Light grey lines represent the variance explained for a single iteration of clustering and fitting. Black line represents the mean variance explained across all iterations. Black bars indicate the percentage change in the variance explained as the number of clusters is reduced. Thick grey line in the testing set indicates the threshold used for determining the optimal number of clusters. e) Spatial patterns of optimal cluster centres (i.e., microstates). f) Statistical significance testing of derived microstates fitted onto the group average data for each stimulus condition. The amount of GFP explained by each microstate at each sample point was tested against the amount of GFP explained by projecting the mean EEG trace under investigation onto baseline data (i.e., the null distribution). g) Microstate projection results on first and second lance group average responses. h) Observed values of differences in total power, onset, and duration (ball-on-stick markers) compared against a non-parametric null distribution of feature differences. Only feature comparisons between microstates which were engaged following both lances could be performed. * Indicates a significant (p≤0.05) difference.

**Figure 3:**
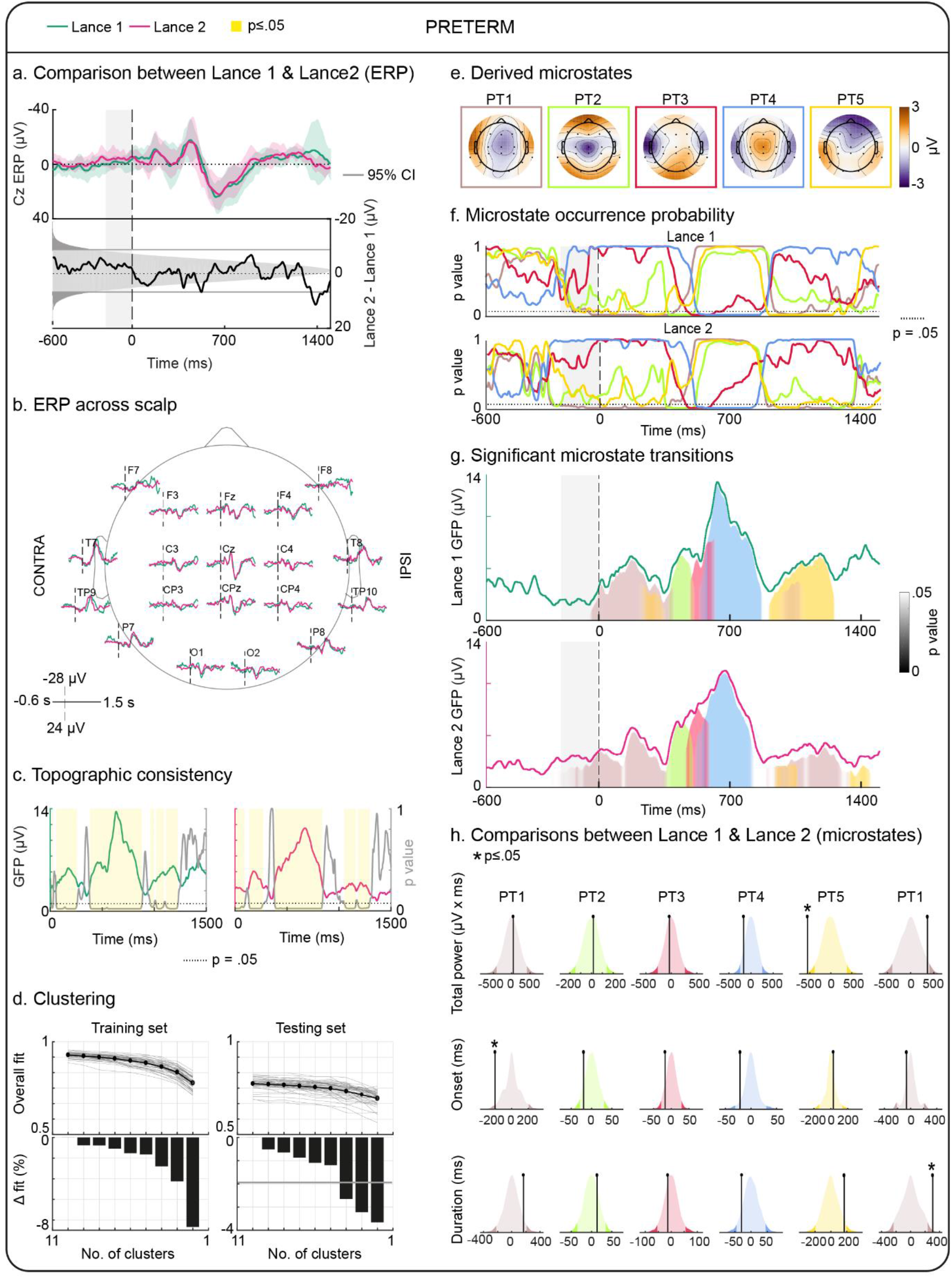
Preterm neonate global event-related field topography (microstate) analysis of EEG data (n=10). See legend for figure 2 for further details.

Changes in single-channel ERPs however can be ambiguous because they can arise from changes in activity magnitude or overall voltage field configuration (Pourtois et al., 2008). To overcome these limitations, we next evaluated cortical responses to the two heel lance procedures using a global event-related field topography (microstate) analysis. Heel lance elicited an increase in global field power (GFP) lasting 1.5 seconds (supplementary figure 2). Figures 2 and 3 show that during this period, first and second lance evoked a consistent pattern of topographies across subjects in both term (figure 2c, first lance: 19-218 and 425-519ms, second lance: 164-244 and 402-498 ms) and preterm (figure 3c, first lance: 58-256 and 382-881 ms, second lance: 134-271 and 382-838 ms) neonates. Topographic consistency across term/preterm neonates was statistically significant (p≤0.05) for 39/63% and 22/61% of the post-lance period following first and second lance respectively. These consistent periods were clustered into six (figure 2d-e) and five (figure 3d-e) microstates for term and preterm infants respectively as this offered the best trade-off between data reduction and data fitting. These number of microstates explained 92/89% of the term/preterm GFP during periods of topographic consistency.

We next asked whether these microstates were engaged at each time point above chance. To do this we compared the projection value of each data point on each of these microstates against a null distribution obtained by calculating the projection of that data point on a random topographic basis obtained from the baseline (−600 – −200 ms) (figures 2f and 3f). At least one microstate (p≤0.05) was assigned to 65/83% and 47/94% of the post-lance period in term/preterm neonates following first and second lance respectively. This analysis also demonstrated transitory stages between microstates, represented by a decaying significance towards the p=0.05 threshold. The output of this analysis is represented in figures 2g (term) and 3g (preterm) which highlights both periods of dominant microstate activity together with transitory periods where multiple states can co-exist.

### Early microstate engagement is reduced in term but not preterm neonates following repeated heel lance

We next examined whether the engagement of microstates differed following first and second lance. In both term and preterm infants, an initial sequence of microstates common to first and second lance was engaged. This consisted of three microstates in term (T1-3) and four in preterm (PT1-4) infants engaged until 700 and 900 ms post-stimulus respectively (figures 2g-h and 3g-h). While there was no significant difference in the engagement of any of these microstates in preterm infants, there was a significant reduction in the engagement of the earliest microstate (T1) in term infants (p=0.016, L01: 410.7 μV×ms, L02: 128.7 μV ×ms). With respect to the timing of engagements, T2 in term infants started significantly later following the second lance (change in onset latency: 135 ms, onset: p=0.002) and PT1 in preterm infants started earlier (change in onset latency: 203 ms, onset: p=0.022).

### Late microstate engagement is altered in both preterm and term neonates following repeated heel lance

The data above showed clear adaptation of short latency microstate activation following a second heel lance in term infants, but not preterm infants. To provide insight into the later phases of cortical processing following heel lance, we next asked whether longer latency microstate engagement changes with repeated stimulation. Neonates of both age groups presented differing late microstate activity following the two heel lances (post 700/900 ms in term/preterm) (figures 2g-h and 3g-h). In term neonates two distinct dominant states (T5, 865-1109 ms, and T6, 1041-1228 ms) were engaged following the first lance and one entirely different microstate (T4, 1025-1306 ms) following the second lance. In preterm neonates, the late cortical microstate pattern following the first heel lance involved the dominant engagement of PT5 (906-1269 ms) and weak engagement of PT1^b^ (955 – 1107 ms). These same microstates were engaged following the second heel lance but with dominant activation of PT1^b^ (892-1349 ms) and weak activation of PT5 (935 – 1068 ms). Both age groups therefore demonstrate in their long latency cortical responses the ability to distinguish and process differentiating components corresponding to each delivered noxious stimulus. This processing appears more refined and specific in developmentally mature term neonates.

### Noxious evoked somatic and autonomic responses habituate upon repeated heel lance in term, but not preterm infants

We next asked whether the somatic and autonomic responses to first and second heel lance, mediated at spinal and brainstem levels, reflected the changes observed at the cortical level. To do this we compared the flexion withdrawal EMG of the stimulated limb, changes in heart rate and facial expression in response to first and second lance.

As expected, noxious stimulation evoked significant changes in biceps femoris muscle activity, heart rate and facial expression scores in both term and preterm infants (p<0.001). Reflex muscle activity increases are between 2.2-4.5 standard deviations of the mean baseline activity in term and 1.3-2.1 in preterm infants (figure 4a-b); median heart rate increases are between 5.4-8.2 beats-per-minute (BPM) from an average resting activity of 132.5 BPM in term and 6.8-9.5 BPM from an average resting activity of 154.6 BPM in preterm infants (figure 4c-d); and median facial expression scores (represented as a percentage change between 0-100%) increased 14.3% in term and between 14.3-71.4% in preterm infants from average baseline scores of 0% (figure 4e-f).

**Figure 4:**
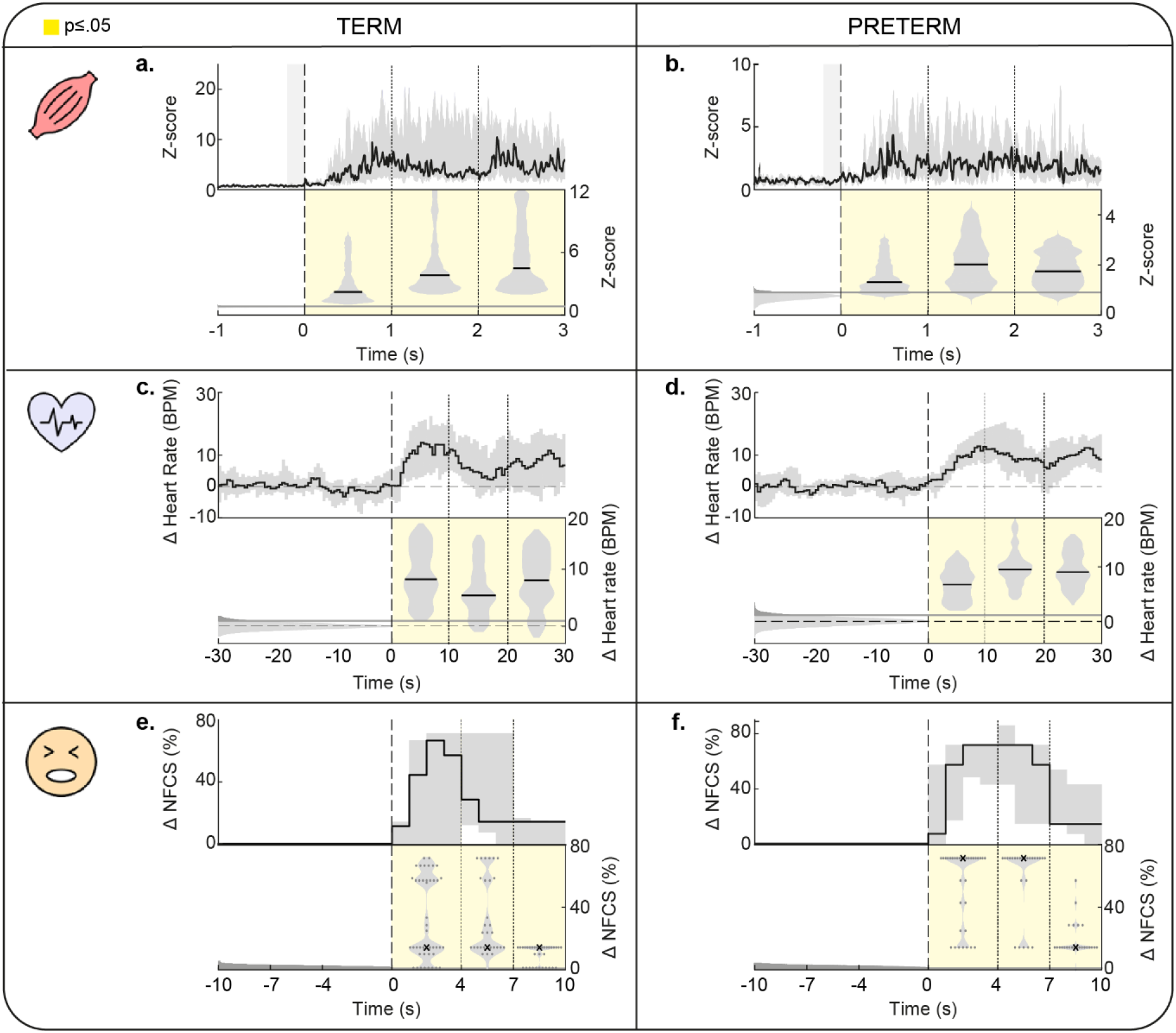
Term (left) and preterm (right) infant somatic and autonomic responses to heel lance. a-b) Changes in the flexion withdrawal reflex of the biceps femoris (EMG) (term n =6, preterm n=4). c-d) Changes in heart rate (derived from ECG) (term n=11, preterm n=10). e-f) Changes in facial expression scores (recorded using video and evaluated using a Neonatal Facial Coding System (NFCS) scale) (term n=11, preterm n=9). Changes in responses were investigated by normalising the data for each measure by the corresponding baseline activity (EMG: −1000 – 0 ms, ECG: −30 – 0 ms, video recording: −10 – 0 ms). Top panels: Solid black trace represents the median response across subjects/trials. Shaded grey area represents the interquartile range. Bottom panels: Solid black lines represent the median response across subjects/trials/time-window. Violin plots represent data distribution within the interquartile range. Median responses within each post-stimulus window were tested against a non-parametric distribution of surrogate baseline medians (shaded grey). The thick grey region represents values above the 95^th^ percentile (grey line). Yellow areas represent time-windows/points with significant changes compared to baseline.

These responses were significantly modulated by noxious stimulus repetition in term, but not preterm infants. In term neonates, the flexion withdrawal reflex changes (1-2 secs post-stimulus, p<0.001), heart rate changes (0-10 secs post-stimulus, p=0.022) and facial expression scores (0-4 seconds post-stimulus, p=0.002) elicited by the noxious stimulus were significantly reduced following the second stimulus compared to the first (figure 5a, c and e). In preterm neonates, these subcortical responses did not change significantly with stimulus repetition (figure 5b, d and f).

**Figure 5:**
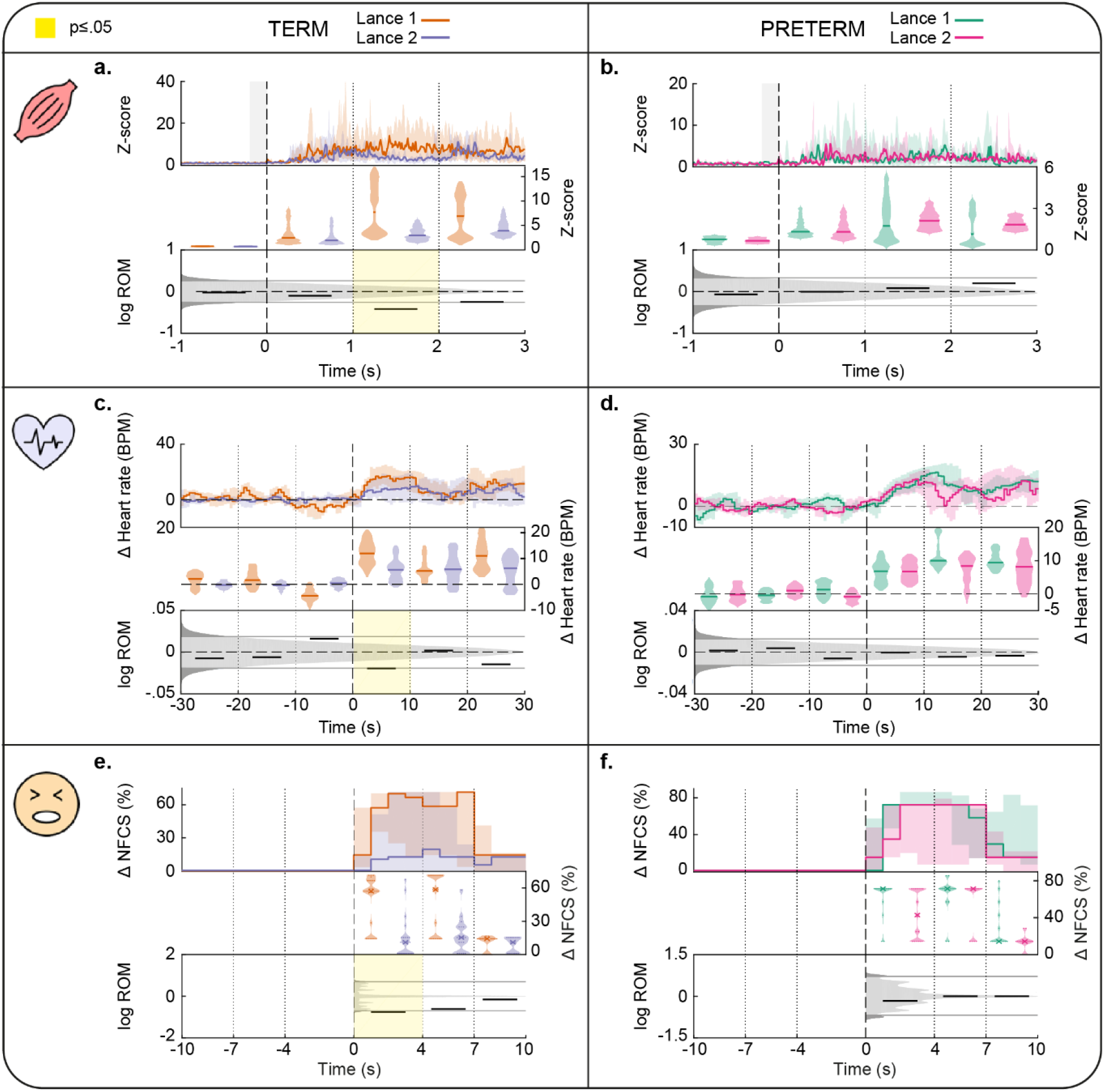
Comparison between first and second lance responses in term (left) and preterm (right) infants. a-b) Differences in the flexion withdrawal reflex of the biceps femoris (EMG) (term n=6, preterm n=4). c-d) Differences in heart rate (derived from ECG) (term n=11, preterm n=10). e-f) Differences in facial expression scores (recorded using video and evaluated using NFCS) (term n=11, preterm n=9). Top panel: Traces represent the median response across subjects. Shaded areas represent the interquartile range at each time-point. Middle panel: Solid lines represent the median response across subjects/temporal window for each lance. Violin plots represent data distribution within the interquartile range. Bottom panel: Solid black lines represent the log ratio of the median (ROM) responses to the two lances within each temporal window. These ratios were tested against a non-parametric distribution of surrogate ratios (shaded grey). Thick grey regions represent values below the 2.5^th^ and above the 97.5^th^ percentiles (thick grey lines). Yellow areas represent time-windows/points with significant differences between first and second lance.

**Figure 6:**
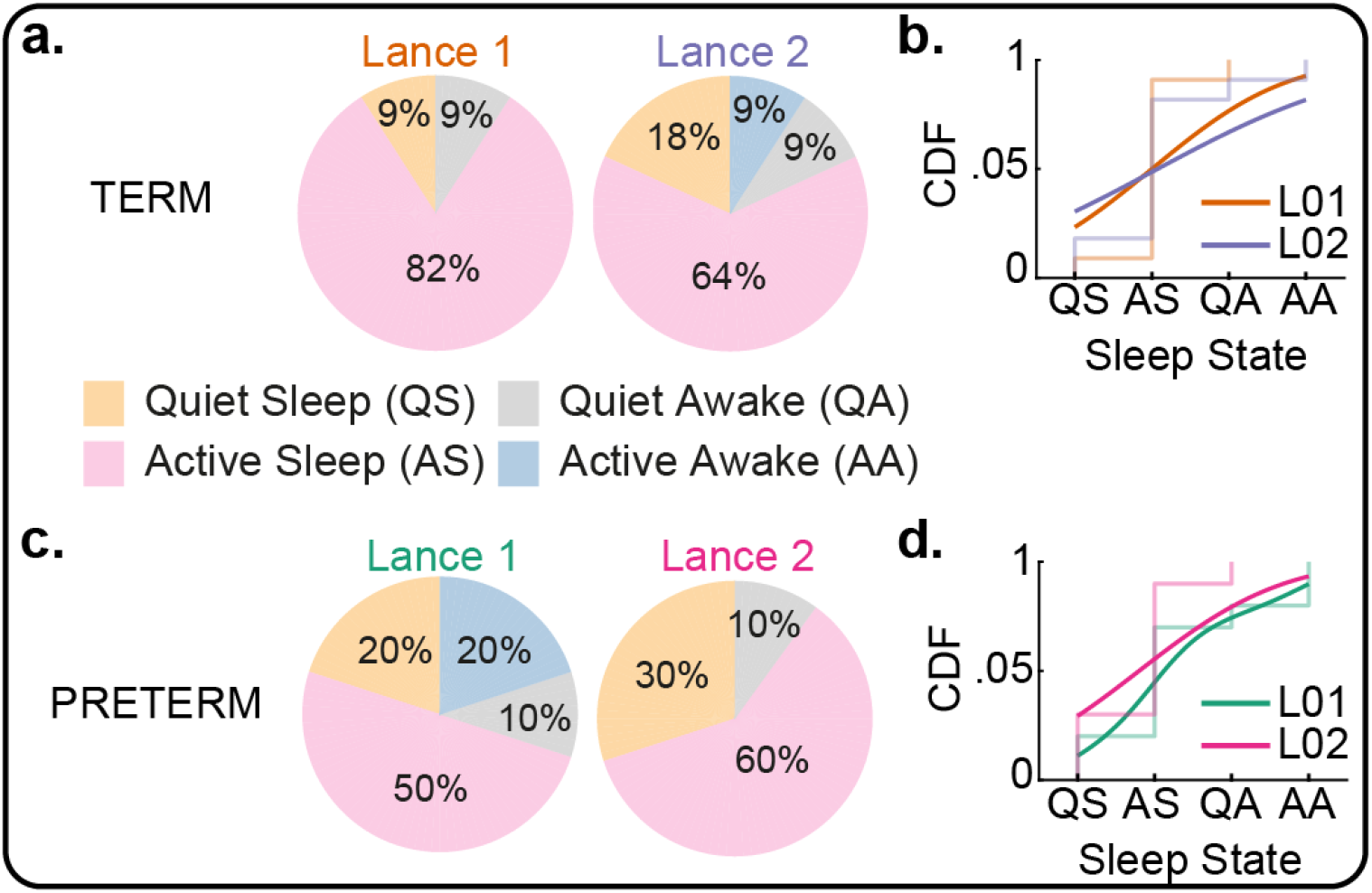
Sleep states distribution immediately prior to the first and second heel lance. Proportion of the 11 term (a) and 10 preterm (c) infants in each of the four sleep states in the immediate 30 seconds prior to the first and second lance. Kolmogorov-Smirnov (K-S) statistical comparison of the two cumulative distributions revealed no significant differences between first and second lance for both term (K-S statistic: 0.09, b) and preterm (K-S statistic: 0.20, d) infants.

## DISCUSSION

Here we have studied the development of habituation to noxious stimulation in newborn infants to test whether the immature human brain can adapt to pain and so encode prediction of imminent nociception. To do this we measured changes in cortical microstates following repeated clinically-required noxious heel lances in term and preterm infants. Microstates are quasi-stable global patterns of scalp potential topographies, and studying their temporal appearance, strength of activation, and cycling behaviour reflects the ongoing simultaneous activity of large-scale pain-related networks. The results show that early microstates, which represent the first tiers of information encoding, were modulated by stimulus repetition in term infants only, while preterm infants did not show any sign of adaptation. Importantly, habituation in physiological (heart rate) and reflex behavioural (facial expression and lower limb reflex withdrawal) responses, which were recorded simultaneously with cortical events also habituated in term, but not preterm infants. Nevertheless, both age groups showed significant changes in the late part of their cortical responses which went beyond simple activity modulation, revealing a complete switch in higher tier cortical processing with repeated identical noxious stimulation. These later processing differences were more distinct in term than preterm infants. These results suggest that the modulatory mechanisms responsible for habituation and altered cortical processing to repeated noxious procedures undergo a developmental shift between preterm and term age.

Our multi-level assessment of responses (cortical, withdrawal reflex, heart rate and behaviour), control for confounds (noxious stimulus site, postnatal age and sleep state (figure 6, methods)), and novel EEG data analysis enabled comparison of responses in cortical and subcortical (autonomic and somatic) circuits to repeated clinically-required noxious stimuli in term and preterm infants. Traditional single channel vertex event-related potential (ERP) analysis has been used extensively to explore responses to noxious stimuli in neonates. This analysis has led to the identification of a specific voltage deflection at the vertex electrode (the P3 potential at Cz) which is associated to the tissue-breaking aspect of the stimulus (Slater, Worley, et al., 2010) and the magnitude of which is associated to factors such as postnatal age (Slater, Fabrizi, et al., 2010), behaviour (Jones et al., 2017) and sex (Verriotis et al., 2018). However, considering the complexity of the pain experience and of the brain networks engaged in the processing of noxious input (Kucyi & Davis, 2015), this approach could be reductive and not allow to capture the multifaceted temporospatial dynamics of nociceptive cortical processing in infants. Alternatively, the topographic information used here has direct neurophysiologic interpretability, in that, pattern differences in time and across conditions (microstates) are indicative of changes in the configuration of multiple active cerebral sources (Michel & Murray, 2012). Moreover, considering EEG microstates as a continuous phenomenon (Mishra et al., 2020) has allowed us to identify periods of dominant single microstate activation, but also others of overlapping multi-microstates co-existence and smooth transitory intervals between dominant microstates. This novel approach to the study of neonatal sensory processing overcomes constrains related to discrete single-channel peak identification and allows for the characterization of a complex continuous dynamic nociceptive processing stream.

The arrival of the nociceptive information to the brain can engage sensory-discriminative and cognitive-affective cortical networks in series or in parallel (Klingner et al., 2016). The short-latency part of the EEG response recorded here is likely to represent the early recruitment of the sensory-discriminative network including the primary somatosensory cortex (SI) and potentially the secondary somatosensory cortex (SII) and part of the insula as this network is the first to be engaged also following innocuous tactile stimulation in both term and preterm neonates (Whitehead et al., 2019). The modulation of these cortical events together with that of the sub-cortical somatic and autonomic responses points to the activation of fast-acting modulatory mechanisms which operate at spinal, brainstem, and cortical level or at spinal level only with effects propagating to higher levels. The shift from non-habituation in preterm to habituation in term infants could therefore reflect maturational changes in these processes, in the mechanisms of their engagement, or both.

In adults, descending pain control involves multiple forebrain regions connecting with the periaqueductal grey region (PAG) and rostral ventromedial medulla (RVM) in the brainstem which regulates nociceptive transmission at the dorsal horn (Millan, 2002). In human term neonates stronger spontaneous pre-stimulus functional connectivity across this descending pain modulatory system predicts a weaker response to a mechanical pin-prick (Goksan et al., 2018). However, this control network is immature in animal models at the equivalent age to human preterm neonates: the descending projections from the brainstem to the dorsal horn are anatomically present, but their inhibitory function is not fully developed (Hathway et al., 2009). A developmental switch subsequently results in the bimodal operation of the RVM, exerting inhibitory controls by regulating GABA and glycine concentration and/or the threshold for C-fibre activation (de Kort et al., 2021; Koch & Fitzgerald, 2014). These maturational changes potentially explain the modulation of the responses in spinally mediated withdrawal reflex, heart rate change, facial behaviours and early cortical events to the second lance procedure observed in term, but not preterm neonates.

A top-down driver to the habituating responses observed could be explained within an associative learning (Dall’Orso et al., 2021) and/or a predictive coding framework (Friston & Kiebel, 2009; Rao & Ballard, 1999) thus requiring information preceding the noxious event to guide subsequent stimulus processing. Predictive-coding models represent the brain as a learning and modelling engine that updates world-related predictions through the encoding of unexpected or novel inputs (i.e., the prediction error) rather than processing all aspects (e.g., saliency, intensity, position, timing) of the input at each occasion (Friston & Kiebel, 2009). According to this model, the changes in microstates following the second lance, may reflect the ability of the brain to predict the qualities of the incoming input common to the first occurrence. In adults, stimulus expectancy affects pain reported and modulates activity in pain related areas such as the anterior cingulate cortex (ACC), insula, but also sensory-discriminative areas such as SI and SII (Atlas & Wager, 2012) which is reflected in event-related potentials amplitude changes (Hird et al., 2018). In particular, the earliest evoked potential components are modulated by the prediction of the incoming stimulus (Bendixen et al., 2012; Rauss et al., 2011), while later responses are more closely coupled with the prediction error (Stefanics et al., 2018). Expectations about a stimuli are normally manipulated before stimulus delivery with instructions to the participants, but they can also develop through basic associative learning processes such as classical conditioning, which is already possible in term neonates (Dall’Orso et al., 2021). The conditioning stimulus in our case (i.e. the cue heralding the arrival of the second lance) could have been holding the lancet against the heel in the 30 seconds before the blade release. This cue could have then allowed the brain to formulate a prediction and compute a prediction error in term, but not preterm infants. Generally, animals act to terminate or avoid pain and responding appropriately to cues that indicate a higher probability of imminent pain makes achieving this goal easier (Fields, 2018). In the absence of other conflicting goals, this would result in the amplification of nociceptive transmission to guide avoidance of potential injury, but this is evolutionary useful only in case actions towards such targets can be taken. As for human neonates where there is no chance to avoid forthcoming injury from the lance, it is likely that the cue signalling imminent pain drives the engagement of the modulatory mechanisms described above which reduces the nociceptive input. These are available in term, but not preterm infants. The fact that preterm neonates do not adapt to repeated unavoidable noxious stimuli could therefore also reflect the immaturity of this cue-prediction system.

Unlike the initial microstate sequence, long-latency microstates were unique following each lance procedure in both preterm and term neonates. These late microstates could represent the activation of the anti-nociceptive (Bingel et al., 2007) or cognitive-evaluative (Petrovic & Ingvar, 2002) networks that modulate subsequent incoming noxious-related signals and/or assess contextual/environmental factors (Tracey & Mantyh, 2007) and are likely to encode the prediction errors in the predictive coding framework (Stefanics et al., 2018). These networks involve the ACC and prefrontal cortex, amongst other high-level brain regions, and together are involved in pain perception and learning processes associated with the prediction or avoidance of noxious stimuli (Coghill et al., 1999; Villemure & Bushnell, 2002). Individual neurons in these high-level brain areas adapt their firing behaviour depending on function (i.e., expectation or prediction error processing) (Dale et al., 2018; Urien et al., 2018), complex organisation of individual brain regions enable multi-processing of stimulus properties, and cortical sites on a global network scale are dynamically coordinated (Geuter et al., 2017). These processes together perhaps allow both long-latency brain activities following the first lance, and brain activities prior to the second lance, to influence the predicted model in anticipation of the second noxious event. In this regard, future work would therefore benefit from studying the cortical microstates pattern prior to each lance event.

The long-latency microstates were more distinct in term compared to preterm neonates for the two lance procedures. In the younger neonate group the late cortical response to the two lances fluctuated closely between two microstates, suggesting potential recycling of a limited number of nodes to integrate complex information from multiple immature cortical sources involved in higher-order stimulus processing (Verriotis et al., 2016). Preterm neonates have several densely connected cortical hubs in place before 30 weeks of gestation but with immature connections (in both length and number) between these hubs regions and the rest of the cortex (Ball et al., 2014; Keunen et al., 2017). These cortico-cortical connections mature over the last trimester of gestation allowing an increase in global network strength and efficiency (Song et al., 2017). This cortical maturation is likely to explain the more distinct microstate patterns observed in term neonates to repeated noxious stimuli. Nevertheless, despite the lack of habituation (i.e., prediction) in preterm neonates, they too demonstrate the potential for high-level processing of contextual differences pertaining to each noxious event, however they seem to be unable to update the levels below. Therefore, their potential inability to relay this processing into the necessary controls that balance threat avoidance and energy wastage may negatively impact the ongoing activity-dependent maturation of cortical circuits and networks. These injury-induced changes to the developing brain are known to result in maladaptive cortical-pain processing and pain-related behaviours associated to negative outcomes later in life or long-lasting disability (Williams & Lascelles, 2020).

This is the first study to demonstrate the implications of closely repeated noxious clinical procedures in human neonates. Term neonates habituate to such stimuli, which appears to arise from differences in stimulus processing at the cortical level and is subsequently reflected by decreased responses across the autonomic and somatic nervous systems. The decreased short-latency responses at the cortical level suggest neonatal learning reduces the processing of nociceptive components common to both events, thus limiting energy wastage. This was subsequently followed by distinct high-level processing of each stimulus. Preterm infants fail to habituate to repeated pain at all levels of their central nervous system, yet immaturely engage neuronal activity patterns suggestive of distinct high-level cortical processing of each noxious event. Given that the preterm architecture is generally geared towards activity-dependent sensory development with low inhibition and high facilitation of inputs, noxious clinical procedures may possibly introduce undesirable and untimely changes, negatively impacting the neonate’s immediate and ongoing developmental trajectory outside of the womb. This could have also impacted the response of our preterm-born neonates studied at term, but numbers were too small to assess this effect.

In conclusion, this work increases our neurophysiological understanding of infant brain responses and behaviours to repeated pain. The data suggest that the preterm brain is capable of encoding high-level contextual differences in pain but cannot update its prediction. The failure of the preterm brain to adapt to repeated pain emphasises the vulnerability of preterm infants to repeated noxious stimulation during their stay in a neonatal intensive care unit.

## METHODS

### Participants

Twenty-one human infants (32–44 completed postmenstrual weeks, 9 female; table 1) who required a clinical blood test and were exposed to two consecutive sampling attempts (with heel lancing) were included in this study. This was an opportunistic sample from a larger database of 283 infants who were recruited from the Maternity and Neonatal Units at University College London Hospitals (UCLH) over 12 years between December 2007 and November 2019. The twenty-one infants included in this study underwent a second heel lance following an unsuccessful first attempt to collect the necessary amount of blood for a clinically-required test and no heel lance was performed solely for the purposes of research. Repeated heel lancing is not common in our units, but is reported elsewhere to occur in 49% of test occasions because of inexperience in conducting the procedure, unsuccessfully reaching the superficial dermal blood vessels, reduced blood flow from the cut, sample haemolysis or an insufficient sample for the desired screening test (Shah & Ohlsson, 2011).

No infants in this cohort were diagnosed with periventricular leukomalacia (PVL), germinal matrix-intraventricular haemorrhage (GM-IVH) greater than grade 2, intrauterine growth restriction (IUGR), trisomy 21, or presented any clinical signs of hypoxic ischaemic encephalopathy (HIE). Ethical approval for this study was given by the NHS Health Research Authority (London – Surrey Borders) and conformed to the standards set by the Declaration of Helsinki. Informed written parental consent was obtained before each study.

### Heel lancing

All heel lances were performed by a trained neonatal nurse using disposable lancets and standard hospital practice. Infants were soothed as and when required. Parents could hold and feed their baby at will. The heel was cleaned with sterile water using sterile gauze and the lancet placed against the heel for at least 30 seconds prior to the release of the blade. This was to obtain a period of recording prior to the stimulus free from other stimulation that could be used as baselines for each of the recording modalities. The heel was squeezed for blood collection only 30 seconds after the release of the blade, again to ensure a post-stimulus period free from other stimuli. The second lance was performed on the same heel due to clinical reasons (i.e., either extensive bruising or damage from previous lances still visible on the other heel) 3-18 minutes (median of 7.5 minutes) after the first (figure 1b). Infants remained in the same position (in skin-to-skin contact with one of their parents, held with clothing or in cot with individualized care) throughout the recording session except for one. Some infants transitioned from one sleep state to another between first and second lance, however there was not a predominant sleep state transition or significant difference in the overall distribution across sleep states for the whole sample (preterm: p=0.975, term: p=1.000) (figure 6).

### Recording set-up

Brain electrical activity at the scalp (electroencephalography, EEG), flexion withdrawal reflex of the lanced leg (surface electromyography, EMG), heart rate changes (electrocardiography, ECG) and facial expressions (video) time-locked to the clinically-required heel lances were recorded (figure 1a). EEG recordings were successful for both lances in all subjects (n=21; 42 recordings). EMG, ECG and video recordings were successful in 26 (n=13), 41 (n=21) and 38 (n=20) recordings respectively.

#### Electroencephalography

The electroencephalogram (EEG) was recorded from a subset of 24 recording electrodes (disposable Ag/AgCl cup electrodes) positioned individually according to the international 10/20 electrode placement system (Homan et al., 1987). Due to partial variability in the electrode montages used over the 12-year study period (i.e., due to the incorporation of multiple study protocols or contraindications), montages were standardised to a 19-electrode setup (figure 1a) that balanced electrode-scalp density with cortical site representation, and minimised interpolating data for a large number of non-recorded channels. The reference electrode was placed at FCz or Fz and the ground electrode at FC1 or FC2 (depending on the position of the infant). Electrode/skin contact impedances were maintained below 10 kOhms when possible, by gently rubbing the skin with a prepping gel (NuPrep, Weaever & Co.) and then applying the electrodes with a conductive paste (10/20 Weaver & Co.). The Neuroscan SynAmps2 EEG/EP (Compumedics, USA) recording system was used to record EEG activity from DC to 500 Hz. Signals were digitized with a sampling rate of 2 kHz and a resolution of 24-bit.

#### Electromyography and Electrocardiography

The electromyogram (EMG) of the lanced leg was recorded from two self-adhesive surface silver/silver-chloride electrodes (Cardinal Health, USA) positioned over the biceps femoris (same ground as that used for EEG recording). The EMG was recorded as a bipolar signal with the same Neuroscan SynAmps2 system used for EEG, amplified (x10,000), and sampled at 2 kHz with a 24-bit resolution.

A lead I electrocardiogram (ECG) was recorded from two self-adhesive surface silver/silver-chloride electrodes (Cardinal Health, USA) positioned over both shoulders (same ground as that used for EEG recording). The ECG was recorded as a bipolar signal with the same Neuroscan SynAmps2 system used for EEG and EMG and sampled at 2 kHz with a 24-bit resolution.

#### Facial actions

Facial expressions were recorded on video and synchronized with the EEG recording with a light emitting diode placed within the frame that was activated by the blade release of the lancet (Worley et al., 2012).

### Data pre-processing

#### EEG data

EEG data was pre-processed using MATLAB (2016, MathWorks, Inc.) and EEGLAB (Delorme & Makeig, 2004). Raw data were filtered with a second-order bidirectional Butterworth bandpass (1–25 Hz) and a notch (48–52 Hz) filter, and epoched between 0.6 s prior to 1.5 s following the stimulus. Raw data were subsequently de-noised using independent component analysis (between 0-3 discrete independent components per trial were removed corresponding to ECG breakthrough or muscle, movement or equipment artifacts, median: 0). Spherical interpolation was then used to estimate data from highly artifactual or non-recorded channels (maximum of four channels per trial, range: 0-4, median: 0). We then applied DC correction, down-sampled the data to 512 Hz and re-referenced to the common average.

#### EMG data

EMG data was pre-processed using custom-written scripts in MATLAB (2016, MathWorks, Inc.). EMG data from 10 out of the 13 subjects was of a quality suitable for analysis. Raw data were filtered with a second-order bidirectional Butterworth bandpass (10-500 Hz) and a notch (48–52 Hz) filter, and epoched between 1 s prior to 3 s following the stimulus. The EMG signal was then converted to z-scores to account for differences in signal amplitude across subjects possibly due to differences in electrode location relative to the biceps femoris and/or contact. The mean and standard deviation for z-scoring were computed from the baseline segments (1 s period prior to the stimulus) for each subject. The z-scored signal was then rectified, and low-pass filtered at 25 Hz (fourth-order bidirectional Butterworth). Artifactual segments were then identified using the median absolute deviation method (Matlab function *‘mad.m’*) within a 1 s sliding window (50% overlap) as data points exceeding three standard deviations from the median signal (Leys et al., 2013). Corrupted segments shorter than 0.05 s were replaced by spline interpolation using 0.05 s worth of data on either side of the corrupted segment, while segments longer than 0.05 s were discarded.

#### ECG data

ECG data was pre-processed using MATLAB (2016, MathWorks, Inc.) and LabChart (version 8, ADInstruments). Raw data were filtered with a second-order bidirectional Butterworth bandpass (0.5 - 45 Hz) filter. Beat-to-beat intervals (R-R intervals) were then automatically detected using the heart rate variability (HRV) module in LabChart and manually confirmed. The resulting signal (heart rate in beats per minute, BPM) was epoched between 30 s prior to and 30 s following the stimulus and baseline corrected using the 30 s period prior to the stimulus.

#### Facial actions

Videos were reviewed offline by a trained behavioural coder. Videos were epoched between 10 s prior to 10 s following the stimulus and infant facial actions were scored second-by-second according to the 7-item version of the Neonatal Facial Coding System (NFCS) (brow bulge, eye squeeze, nasolabial furrow, open lips, vertical stretch mouth, horizontal stretch mouth, and taut tongue) (Ahola Kohut & Pillai Riddell, 2009; Craig et al., 1993; Grunau & Craig, 1987). The facial features were scored as either present (1) or not present (0) resulting in a percentage total score at each second (where a score of 100% indicates that all 7 facial actions were observed). Where the view of infants facial actions were partially/fully obstructed, data were estimated with conservative judgements based on either the assumption of facial symmetry, cry and body movements, and behaviours either side of the missing period (Ahola Kohut & Pillai Riddell, 2009; Pillai Riddell et al., 2007). Partial estimates of facial scores were made in 5 out of the 38 recordings.

### Analysis of responses to the lance

We then assessed whether a lance stimulus elicited significant changes in EEG, EMG, ECG and facial actions in preterm and term infants separately. This analysis was conducted by pooling data from first and second lance attempts and comparing the signal following stimulation against baseline. Due to the high variability in neonate EMG, ECG and facial action responses to the noxious stimuli, the median summary statistic was analysed for these measures as a method of more robustly capturing the central tendency for each dataset.

#### EEG

EEG changes were assessed as traditional single channel event-related potentials, and as global field power changes.

##### Single channel event-related potential (ERP)

The group ERP waveform was first estimated by averaging individual recordings at Cz across subjects/trials. Significant changes from baseline following stimulation were determined with sample-by-sample non-parametric testing against a null distribution obtained from baseline data. This was the distribution of the baseline samples of an array of 5000 grand averages obtained from 5000 phase-randomised versions of the collected dataset. Such surrogate data were derived based on the assumption of temporal stationarity of variation around the mean baseline signal across subjects/trials (see section *‘Surrogate data generation’* for further details). To account for multiple testing, only continuous periods of samples outside the 95% confidence interval (CI) that lasted for longer than 5% of the post-lance period were considered significant (Guthrie & Buchwald, 1991).

##### Global Field Power (GFP)

The group GFP was estimated by averaging individual recordings at each channel across subjects/trials and then computing the standard deviation across the channel dimension. Significant changes from baseline following stimulation were determined with sample-by-sample non-parametric testing against a null distribution obtained from baseline data. This was the distribution of the baseline GFP samples of an array of 5000 grand averages obtained from 5000 phase-randomised versions of the collected dataset (see ‘*Surrogate data generation*’ for further details). To account for multiple testing, only continuous periods of samples outside the 95% CI that lasted for longer than 5% of the post-lance period were considered significant (Guthrie & Buchwald, 1991).

#### EMG

The median signal across subjects/trials/temporal window was determined for three 1 -second-long segments post-stimulus and compared against a non-parametric null distribution obtained from baseline data. This was the distribution of the median value across subjects/trials/temporal window of 1-second-long baseline segments from 5000 phase-randomised versions of the collected dataset (see *‘Surrogate data generation’* for further details). Phase randomisation was performed on the bandpass filtered baseline EMG signal before signal rectification. Post-stimulus changes were considered significant if the median response was outside the 95% CI of this null distribution.

#### ECG

The median signal across subjects/trials/temporal window was determined for three 10-second-long segments post-stimulus and compared against a null distribution obtained from baseline data. This was the distribution of the median value across subjects/trials/temporal window of 10-second-long baseline segments from 5000 phase-randomised versions of the collected dataset (see *‘Surrogate data generation’* for further details). Phase randomisation was performed on the processed heart rate signal. Post-stimulus changes were considered significant if the median response was outside the 95% CI of this null distribution.

#### Facial actions

The median signal across subjects/trials/temporal window was determined for two 3-second-long and one 4-second-long segments post-stimulus and compared against a null distribution obtained from baseline data. This was the distribution of the median value across subjects/trials/temporal window of two 3-second-long and one 4-second-long baseline segments from 5000 datasets obtained by bootstrapping 78 lance video-recordings (68 of which were not part of this study and a total of 41 video-recordings were those of preterm neonates). Post-stimulus changes were considered significant if the median response was outside the 95% CI of this null distribution.

### Analysis of response differences to repeated noxious stimuli

We finally assessed differences in responses following first and second lance in EEG, EMG, ECG and facial actions in preterm and term infants separately.

#### EEG

Differences in responses to first and second lance were determined as differences in single channel event-related potential and in global event-related field topography (microstates).

##### Single channel event-related potential

The group ERP waveform was first estimated by averaging individual recordings at Cz across subjects for each stimulus condition. Significant differences between first and second lance responses were determined at each sample by comparing differences against a non-parametric null distribution. This was the distribution of the same difference between an array of 5000 surrogate pairs of group averages obtained by phase randomising and resampling the collected datasets (see *‘Surrogate data generation’* for further details). To account for multiple testing, only continuous periods of samples outside the 95% CI that lasted for longer than 5% of the post-lance period were considered significant (Guthrie & Buchwald, 1991).

##### Global event-related field topography

Differences in global event-related field topography were determined with microstate analysis using Ragu (Habermann et al., 2018; Koenig et al., 2011) and custom-written MATLAB scripts. Microstates are scalp potential fields which maintain a semi-stable topography over transient periods of 60-120 ms (Khanna et al., 2015; Michel & Koenig, 2018). Microstate activation and switching across time represent changes in brain network engagement and information transfer (Custo et al., 2017). In this study, we considered microstates as a continuous phenomenon and took a geometric approach similar to that described in (Mishra et al., 2020), where each microstate forms the vector basis of a reduced subspace of the original channel space. With this approach individual time points can be labelled as multiple microstates and microstate transitions are gradual rather than discrete.

We first determined our microstate basis. This was achieved by firstly identifying periods of consistent topography across subjects/trials following noxious stimulation with a topographic consistency test. This test assumes that if similar cortical sources are engaged in response to the same stimulus, the resulting event-related potential field would appear as a consistent spatial distribution across trials (and subjects) (Koenig & Melie-García, 2010). Data which were topographically consistent across trials/subjects were then pooled in the same channel space and clustered to define our microstate basis. Clustering was performed with the aim to identify distinct topographies representing different cortical source configurations (in both location and/or orientation) that account for most of the variance in the data. Clustering was performed using a modified hierarchical algorithm to that provided in the Ragu toolbox (Habermann et al., 2018):

1. As part of a cross-validation process, the full dataset was randomly split into a training set (50%) and a test set (50%).
2. In the training set each sample was initially considered as the centre of its own distinct cluster.
3. Pair-wise spatial correlation between cluster centres was calculated leading to a covariance matrix.
4. Pairs of clusters whose centres were maximally correlated were merged and a new cluster centre was calculated as the average of the two original cluster centres. These formed the current template maps.
5. The explained variance in the test set was calculated for the template maps.
6. Steps 3 to 6 were repeated until we were left with two clusters. This led to an explained variance versus number of clusters relationship for a single cross-validation iteration.
7. 50 cross-validation iterations were repeated.
8. The 50 explained variance versus number of cluster relationships were averaged and the percentage change in the average variance explained between neighbouring cluster sets was calculated.
9. The optimal number of microstates to use was selected as the size of the last cluster set before the average variance explained dropped by more than 2% with another aggregation step. This choice represents a parsimonious use of microstates that still explain a considerable fraction of signal energy (in this case ≥80% of the GFP signal) (Murray et al., 2008).
10. The final microstate basis was determined on the full dataset.

Once the microstate basis was defined, we calculated the projection of each data point of the first and second lance group average signal on all microstates (Mishra et al., 2020).

Finally, to understand whether the proximity of a data point to a microstate was above chance (i.e., whether a given microstate was indeed engaged at that time), we compared the projection value of the data point on each microstate against a non-parametric null distribution. This was obtained by calculating the projection of that data point on a random topographic basis which consisted of the baseline topographies of all trials (subjects × conditions) in the dataset. In order to consider the microstate significantly engaged, the projection value had to be above the 95% percentile of this null distribution, and the duration of significant engagement had to continue for longer than 5% of the post-lance period (Guthrie & Buchwald, 1991). The periods in which microstates were significantly engaged were called *microstate occurrences*. Distinct occurrences of the same microstate were considered to be separated if the period in between was longer than the mean duration of occurrence across both lances. Microstate occurrences that were separated by a duration shorter than this value were considered to be part of the same event.

We then calculated the onset, duration and variance explained (total power) for each microstate occurrence for first and second lance attempts. Onset was the first time-point in which the projection value exceeded the significance threshold, duration was the difference between onset-offset (where offset is the last time-point in which the projection value exceeded the significance threshold) and variance explained (total power) was the overall amount of variance explained across the whole occurrence duration. These metrics were then compared between first and second lance. This was accomplished by comparing the observed difference in these metrics against a non-parametric null distribution. This was the distribution of the differences in these metrics calculated from an array of 5000 surrogate pairs of group averages obtained by phase randomising and resampling the collected dataset (see *‘Surrogate data generation’* for further details). Metric differences outside the 95% CI of this null distribution were considered significant.

#### EMG

Significant differences between first and second lance were determined for three 1-second-long segments post-stimulus with non-parametric testing. The log ratio between the median signal across subjects/temporal window for all 1-second-long segments following first and second lance (baseline and post-stimulus periods) were calculated and compared against a null distribution. This was the distribution of the same ratio between an array of 5000 pairs of surrogate groups obtained by phase randomising and resampling the collected dataset (see *‘Surrogate data generation’* for further details). Phase randomisation was performed on the bandpass filtered EMG signal before signal rectification. Differences in post-stimulus temporal windows were considered significant if the observed log ratio of the median responses was outside the 95% CI of this null distribution.

#### ECG

Significant differences between first and second lance were determined for three 10-second-long segments post-stimulus with non-parametric testing. The log ratio between the median signal across subjects/temporal window for all 10-second-long segments following first and second lance (baseline and post-stimulus periods) were calculated and compared against a null distribution. This was the distribution of the same ratio between an array of 5000 pairs of surrogate groups obtained by phase randomising and resampling the collected dataset (see *‘Surrogate data generation’* for further details). Phase randomisation was performed directly on the processed heart rate signals. Differences in post-stimulus temporal windows were considered significant if the observed log ratio of the median responses was outside the 95% CI of this null distribution.

#### Facial actions

Significant differences between first and second lance were determined for two 3-second-long segments and one 4-second-long segment post-stimulus with non-parametric testing. The log ratio between the median signal across subjects/temporal window for these three segments following first and second lance were calculated and compared against a null distribution. This was the distribution of the same ratio between an array of 5000 pairs of groups obtained by resampling 78 lance video-recordings (68 of which were not included in this study). Differences in post-stimulus temporal windows were considered significant if the observed log ratio of the median responses was outside the 95% CI of this null distribution.

### Surrogate data generation

Phase randomisation is a method that enables surrogate data to be generated for non-parametric statistical hypothesis testing (Theiler et al., 1992). Surrogate data can be used to form a null distribution for significance testing of original data as they can be considered as random copies of non-linear time series that preserve physiological correlations through the preservation of the signals power spectrum (Prichard & Theiler, 1994).

Each recording time-locked to a stimulus can be considered the linear sum of a signal of interest (i.e., the response to a stimulus) and a stationary random noise component. Based on the assumption that the signal is the same in each recording while the noise changes, conducting another recording should result in the data being a linear sum of the same signal but with different random noise. Creating surrogate data therefore consists of generating new random noise to add to the signal estimated from the recorded data. Surrogate data generation begins by isolating the noise from the signal by removing the group average (i.e., the non-stationary model component) from each subject’s recording. This is not necessary when generating surrogate baseline segments as the recorded baseline is assumed to just represent stationary random noise with no signal. Each noise time-series is then converted into the frequency domain and the phase at each frequency is rotated by an independent random value between 0 and 2π (Theiler et al., 1992). Applying an inverse Fourier transformation leads to a new time series which has the same spectral characteristics of the original noise, but a different temporal realisation (Theiler & Prichard, 1996). Finally, adding back the estimated signal (i.e., the group average or 0 for baseline) results in a new surrogate dataset, which can be used to generate a sample of the null non-parametric distribution of interest.

When comparing parameters between first and second lance the signal subtracted from the data to isolate the noise is estimated for first and second lance separately. However, to prevent the introduction of systematic differences between surrogate data when establishing the non-parametric null distribution, the signal added back at the end of the surrogate data generation process is the grand average across both lances. The surrogate time-series are then pooled together, and two new groups are formed with resampling. Differences between these two groups are a sample of the null-distribution of interest.

## Acknowledgments

The research was performed at the University College London Hospitals (UCLH) Maternity and Neonatal Units. The authors thank the families of the infants that participated in this research. The authors thank Stephanie Koch for discussions of the results.

## SUPPLEMENTARY DATA

**Supplementary Figure 1:**
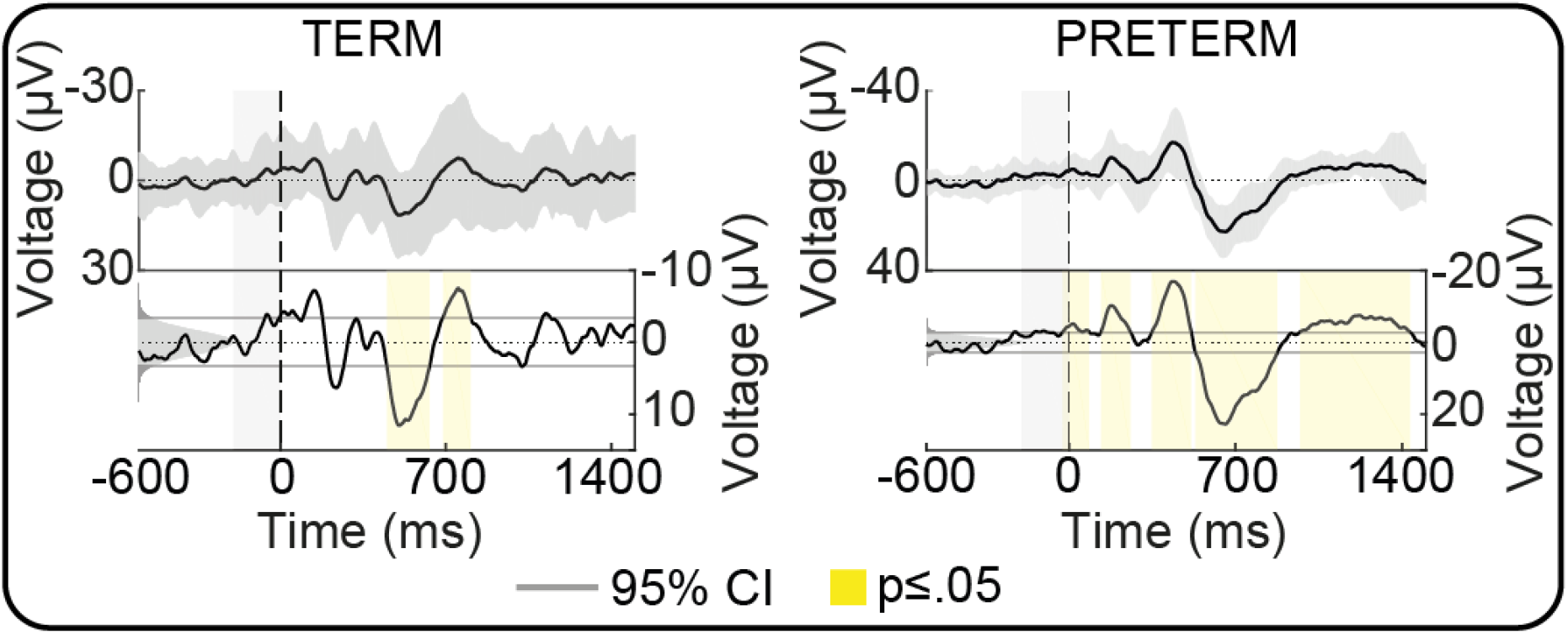
Vertex (Cz) event-related potential of the mean activity across subjects/trials in each infant group (left, term infants; right, preterm infants) in response to heel lancing. Top panels: mean (black line) and standard deviation (shaded grey area) voltage across subjects/trials. Bottom panels: non-parametric distribution of surrogate baseline means (shaded grey) and values below the 2.5^th^ and above the 97.5^th^ percentiles (thick grey regions and lines). Yellow areas represent time-periods with significant changes compared to baseline. The results indicate significant changes in the voltage potential at the vertex post-stimulus. A greater period of the post-lance response is significant in preterm than term neonates.

**Supplementary Figure 2:**
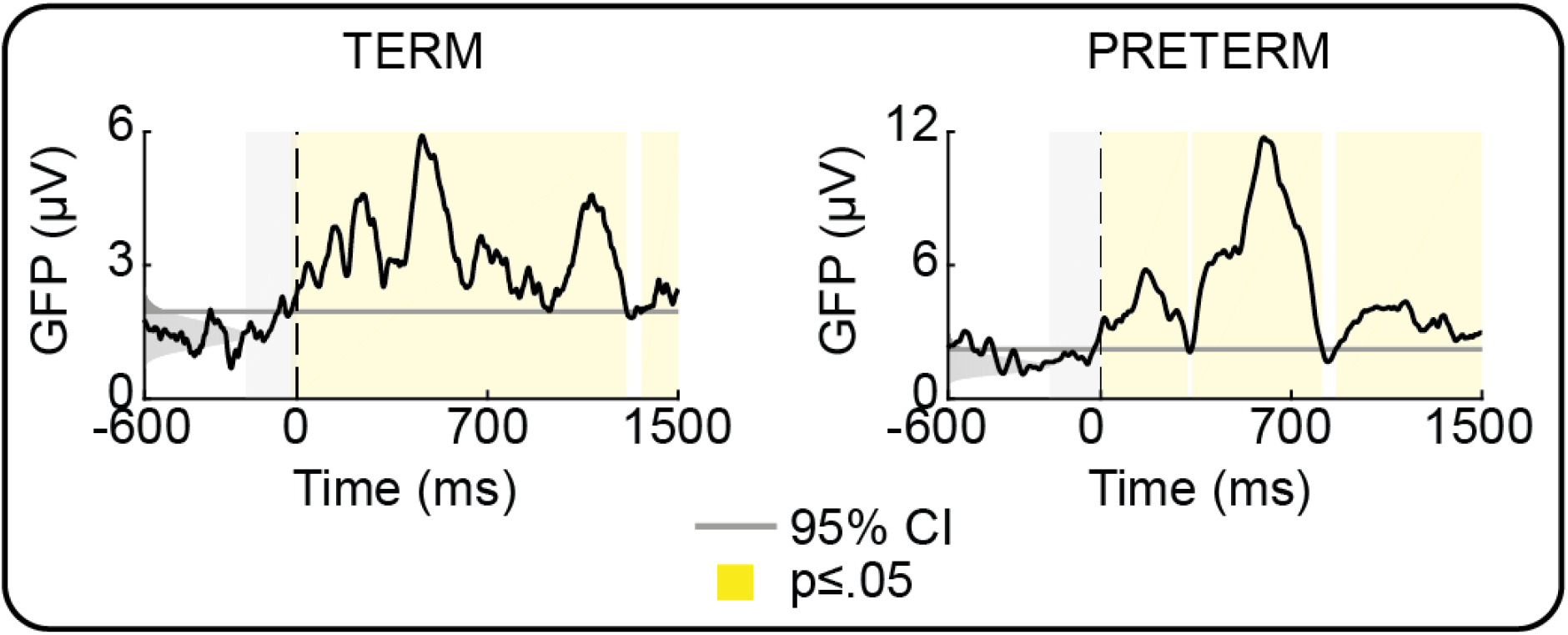
Global field power (GFP, i.e., the spatial standard deviation) of the mean scalp activity across subjects/trials in each infant group (left, term infants; right, preterm infants) in response to heel lancing. The observed GFP response at each sample point was tested against a non-parametric distribution of surrogate baseline GFPs (shaded grey). The thick grey region represents values above the 95^th^ percentile (grey line). Yellow areas represent time-periods with significant changes compared to baseline. The results indicate significantly increased cortical spatial variance following the noxious procedure by both preterm and term neonates.

## Notes

**Conflict of Interest Statement**, The authors declare no conflict of interest in this study.

### Competing Interest Statement

The authors have declared no competing interest.

